# Unique retinal binding pocket of primate blue-sensitive visual pigment

**DOI:** 10.1101/2020.05.29.123463

**Authors:** Yuki Nonaka, Shunpei Hanai, Kota Katayama, Hiroo Imai, Hideki Kandori

## Abstract

The visual pigments of humans contain 11-*cis* retinal as the chromophore of light perception, and its photoisomerization to the all-*trans* form initiates visual excitation in our eyes. It is well known that three isomeric states of retinal (11-*cis*, all-*trans*, and 9-*cis*) are in photoequilibrium at very low temperatures such as 77 K. Here we report the lack of formation of the 9-*cis* form in monkey blue (MB) at 77 K, as revealed by light-induced difference FTIR spectroscopy. This indicates that the chromophore binding pocket of MB does not accommodate the 9-*cis* form, even though it accommodates the all-*trans* form by twisting the chromophore. Mutation of the blue-specific tyrosine at position 265 into tryptophan, which is highly conserved in other animal rhodopsins, led to formation of the 9-*cis* form in MB, suggesting that Y265 is one of the determinants of the unique photochemistry in blue pigments. We also found that 9-*cis* retinal does not bind to MB opsin, implying that the chromophore binding pocket does not accommodate the 9-*cis* form at physiological temperature. The unique property of MB is discussed based on the present results.

---

Humans have two kinds of vision: twilight vision mediated by rhodopsin in rod photoreceptor cells and color vision achieved by multiple color pigments in cone photoreceptor cells.^1^ Humans also possess three color pigments, red-, green-, and blue-sensitive proteins, that maximally absorb at 560, 530, and 425 nm, respectively.^2^ Rhodopsin and color pigments both contain a common chromophore molecule, 11-*cis* retinal, and different chromophore-protein interactions allow preferential absorption of different colors.^3,4^ Purified proteins are a prerequisite to understanding the mechanism of color tuning and the activation process by light. Studying rhodopsin is highly advantageous because large amounts of protein can be obtained from vertebrate and invertebrate native cells. Bovine rhodopsin is a standard protein, and its crystal structures have been reported for the unphotolyzed state,^5^ opsin,^6^ photobleaching intermediates,^7,8^ active state,^9,10^ and the active-state complexed with the C-terminus peptide of the α subunit of G-protein,^10–12^ or engineered mini-Go,^13^ and arrestin.^14^ These structures have provided insight into the mechanism of the chromophore-protein interaction and activation.

On the other hand, structural studies of color pigments lag far behind those of rhodopsin. No color visual pigments have yet been crystallized. Under such circumstances, we started structural studies of primate color pigments by using low-temperature difference FTIR spectroscopy.^15,16^ To achieve this, monkey red- (MR), and monkey green- (MG)-sensitive color visual pigments were expressed in HEK cells, and light-induced difference FTIR spectra were measured at 77 K, where the photoproduct, batho-intermediate (Batho), was reverted to the initial state by light.^17–21^ Consequently, photoconversions from the initial state to Batho, and Batho to the initial state were repeated. This photochromic property has been highly advantageous to improve the signal-to-noise ratio of FTIR spectra. Although a lower expression level was reported for blue-sensitive pigments, we successfully obtained light-induced difference FTIR spectroscopy of MB from marmoset.^22^

In the study of MR and MG, analysis of the 9-*cis* form was useful. In the study of bovine rhodopsin, it is well known that the 11-*cis* form (the unphotolyzed state), the all-*trans* form (Batho), and the 9-*cis* form (also called isorhodopsin) are in photoequilibrium at 77 K,^23,24^ indicating that the retinal binding pocket of bovine rhodopsin accommodates these three isomers. This is also the case for other visual pigments including MR and MG.^17^ In fact, a structural comparison between the 11-*cis* and 9-*cis* forms provided useful information of protein-bound water molecules in MR and MG. We reported that water O-D stretching vibrations in D_2_O are identical in both 11-*cis* and 9-*cis* forms for bovine rhodopsin,^24^ suggesting the same hydrogen-bonding networks. In contrast, in MR and MG, we found different water O-D stretching vibrations between the 11-*cis* and 9-*cis* forms, leading to positional identification of these protein-bound water molecules.^18^ When we published the first paper on MB, we focused on the difference spectra between the 11-*cis* (unphotolyzed state) and *all*-trans (Batho) forms.^22^

In the present study, we attempted to stabilize the 9-*cis* form of MB at 77 K. Initially UV-visible spectroscopy was used to study the photoequilibrium mixture at liquid air or nitrogen temperatures.^23,25–28^ However, vibrational spectroscopy such as resonance Raman and IR spectroscopy is more suitable for this aim, as isomer-specific vibrations appear in the 1250-1150 cm^−1^ region, and since each isomer can be easily distinguished.^24,29–35^ Interestingly, when we illuminated MB at various wavelengths of light in this study, we could not observe the 9-*cis* form of MB at 77 K. This indicates that the chromophore binding pocket of MB does not accommodate the 9-*cis* form, even though it does accommodate the all-*trans* form (Batho). Mutation of the blue-specific tyrosine at position 265 into tryptophan, which is highly conserved in other animal rhodopsins, led to formation of the 9-*cis* form in MB, suggesting that Y265 is the determinant of the unique photochemistry in blue pigments. We also found that 9-*cis* retinal did not bind to MB opsin, implying that the chromophore binding pocket does not accommodate the 9-*cis* form. The unique property of MB is discussed based on the present results.

## MATERIALS AND METHODS

### Sample Preparation

MB cDNA was tagged with the Rho1D4 epitope sequence and introduced into the pcDNA3.1 expression vector. This construct was expressed in the HEK cell line and regenerated with 11-*cis*-retinal.^22^ MRh was prepared similarly.^22^ Site-directed mutagenesis was performed using the QuikChange Multisite-Directed Mutagenesis Kit (Agilent Technologies, Inc., Santa Clara, CA, USA).^22^ The regenerated sample was solubilized with a buffer containing 2% (w/v) *n*-dodecyl-β-D-maltoside (DDM) (final concentration was 1% (w/v)), 50 mM HEPES, 140 mM NaCl, and 3 mM MgCl_2_ (pH 7.0) and purified by adsorption on an antibody-conjugated column and eluted with a buffer containing 0.10 mg/mL 1D4 peptide, 0.02% DDM, 50 mM HEPES, 140 mM NaCl, and 3 mM MgCl_2_ (pH 7.0).

### Low-Temperature FTIR Spectroscopy

For FTIR spectroscopy, MB and MRh samples in detergent were reconstituted into phosphatidylcholine (PC) liposomes with a protein-to-lipid molar ratio of 1:30 by dialysis to remove DDM. The reconstituted sample was suspended in a buffer containing 2 mM phosphate and 10 mM NaCl (pH 7.25), placed onto a BaF_2_ window and dried with an aspirator. Low-temperature FTIR spectroscopy was applied to the films hydrated with H_2_O at 77 K, as described previously.^17,18,22^

The MB samples were illuminated with 400 nm light (by using an interference filter) for 5 min at 77 K, followed by illumination with >520 nm light (by using a VO54 cut-off filter) for 5 min at 77 K.^22^ The former and latter illuminations convert MB to Batho, and Batho to MB, respectively, from which Batho (all-*trans*) minus MB (11-*cis*) difference FTIR spectra were obtained. To examine the 9-*cis* form, Batho was illuminated by using various cut-off filters such as VY47 (>450 nm light), VY46 (>440 nm light), and VY45 (>430 nm light). For each measurement, 128 interferograms were accumulated, and 40 recordings were averaged. FTIR spectra were recorded with a 2 cm^−1^ resolution.

### Hydroxylamine Bleach

HEK cells expressing each protein were divided into two, to which either 11-*cis* or 9-*cis* retinal was added and incubated for 1 hour. Then, the regenerated sample was solubilized with a buffer containing 2% (w/v) DDM (final concentration was 1% (w/v)), 50 mM HEPES, 140 mM NaCl, and 3 mM MgCl_2_ (pH 7.0). The DDM-solubilized proteins in the presence of 10 mM hydroxylamine were kept for 30 min in the dark at 4°C, and then illuminated with >400 nm (the wild-type and mutant MB) or >480 nm (the wild-type and mutant MRh) light for 1.5 min, respectively.

## RESULTS

### No Formation of the 9-cis Form in MB at 77 K

The dotted lines in Figure 2a show light-minus-dark difference FTIR spectra upon formation of Batho from MB, MRh, MG, and MR, where positive and negative signals correspond to all-*trans* and 11-*cis* retinal, respectively. In these cases, Batho was formed by illumination of a monochromatic light from an interference filter (400 nm for MB, 501 nm for MRh and MG, and 543 nm for MR).^22^ On the other hand, photo-reversion was possible by illumination with light at longer wavelengths derived from a cut-off filter (>520 nm for MB, >610 nm for MRh and MG, and >660 nm for MR), which was evidenced by mirror-imaged difference spectra in the UV-visible or IR regions.^22^

**Figure 1:**
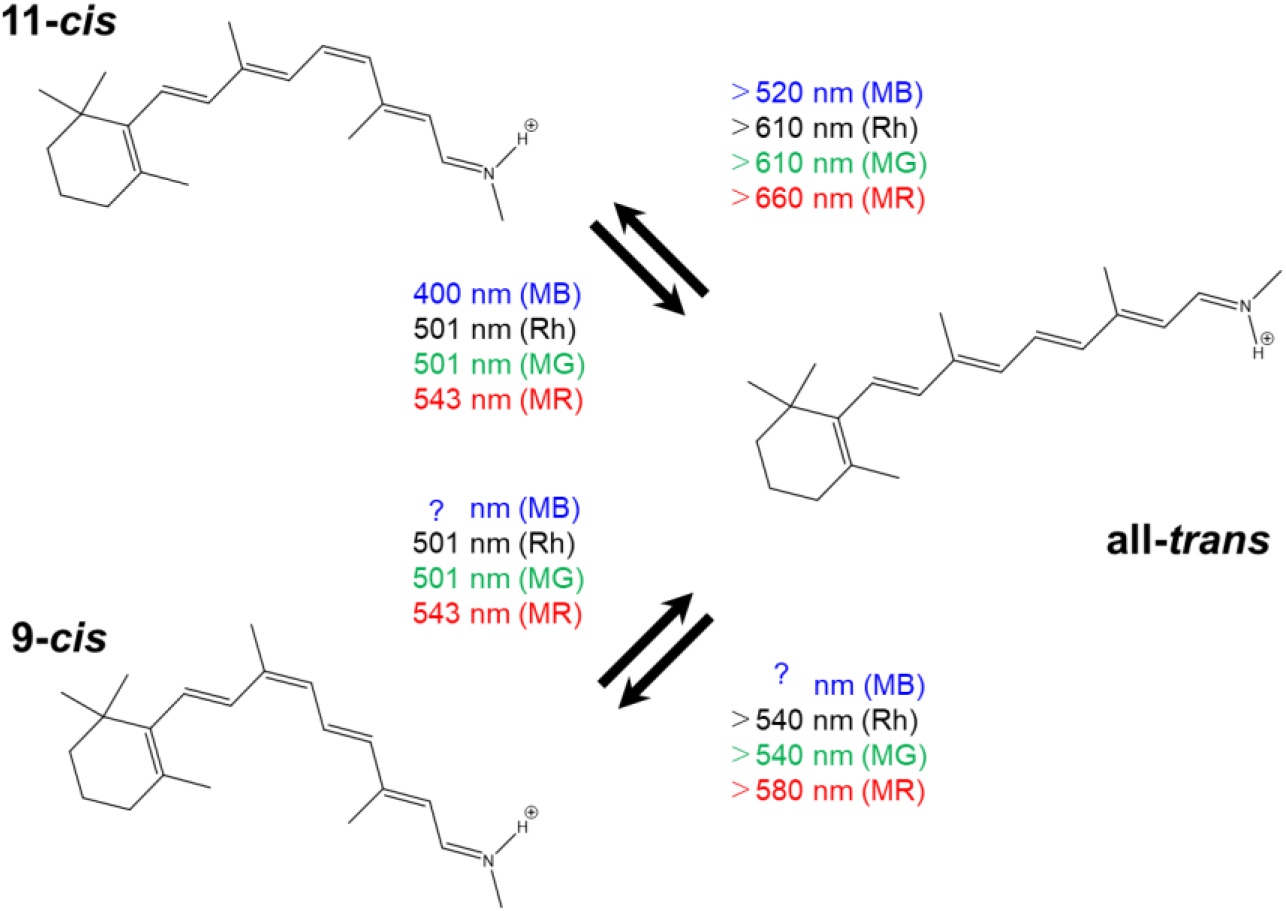
Schematic drawing of the photoequilibria in MR, MG, MB, and MRh. The unphotolyzed state contains the 11-*cis* form, while illumination leads to photoequilibria mainly containing the all-*trans* form (Batho-intermediate) or the 9-*cis* form. Illumination wavelengths were established for MR MG, and MRh. On the other hand, photoequilibria were only investigated between the 11-*cis* and all-*trans* forms for MB.^22^

**Figure 2:**
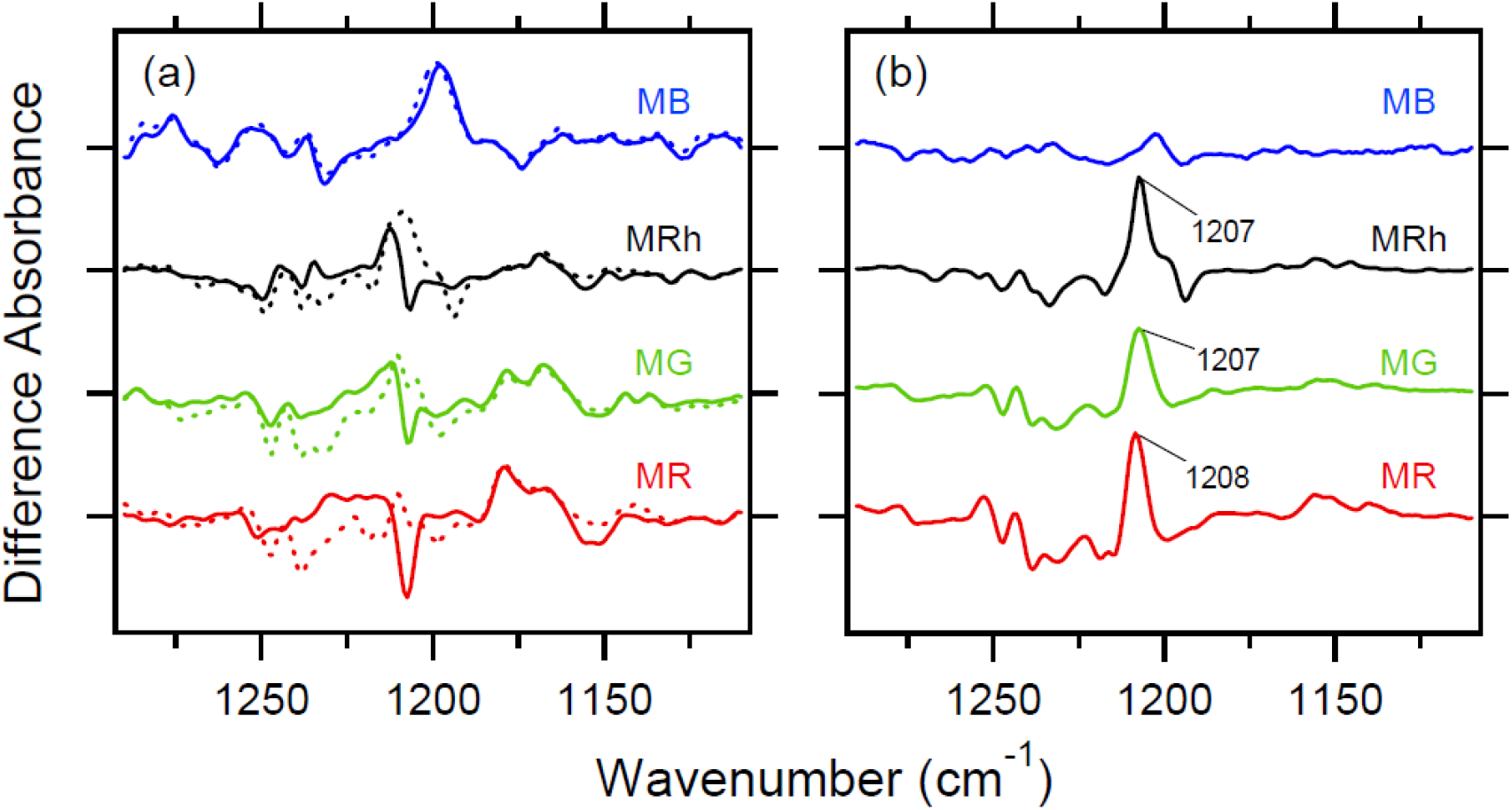
(a) Light-minus-dark difference FTIR spectra in the 1300-1100 cm^−1^ region for MB, MRh, MG, and MR at 77 K. Dotted lines were obtained by the repeated illuminations of 400 nm and >520 nm lights for MB (blue line), 501 nm and >610 nm lights for MRh (black line) and MG (green line), and 543 nm and >660 nm lights for MR (red line). Solid lines were obtained by the repeated illuminations of 400 nm and >430 nm lights for MB (blue line), 501 nm and >540 nm lights for MRh (black line) and MG (green line), and 543 nm and >580 nm lights for MR (red line). In the case of MRh, MG, and MR, dotted and solid lines correspond to the all-*trans* minus 11-*cis*, and all-*trans* minus 9-*cis* difference spectra, respectively. (b) Double difference spectra in (a), where the solid line was subtracted from the dotted line. In the case of MRh, MG, and MR, this corresponds to the 9-*cis* minus 11-*cis* difference spectra.

While the dotted lines in Figure 2a originate only from all-*trans* and 11-*cis* forms, illumination of Batho with different wavelengths yields formation of the 9-*cis* form. For instance, the solid black line in Figure 2a represents the all-*trans* (Batho) minus 9-*cis* spectrum for MRh in which Batho was formed by illumination at 501 nm, whereas the illumination of Batho with >540-nm light accumulated the 9-*cis* form. The reason is that the 9-*cis* form has the most blue-shifted absorption, and its marker band is a peak at 1207 cm^−1^.^24,30,35^ The black spectrum in Figure 2b is the double difference of the black dotted and solid spectra in Figure 2a, where positive and negative signals correspond to 9-*cis* and 11-*cis* retinal, respectively. This is also the case for the reported color visual pigments. To accumulate the 9-*cis* form, we established illumination wavelengths as >540 nm and >580 nm for MG and MR, respectively,^18^ and the 9-*cis*-specific vibrational band at 1207 cm^−1^ (1208 cm^−1^ for MR) was used to optimize the experimental conditions.

When converting wavelengths from Batho into 9-*cis* and 11-*cis* forms, the difference was 70 nm for MRh and MG, and 80 nm for MR. In the present study, we attempted to accumulate the 9-*cis* form for MB similarly at 77 K. However, we found an unusual photochemical property for MB. The solid blue line in Figure 2a is the difference FTIR spectrum obtained by illuminating Batho at >430 nm for MB, which coincides with the spectrum obtained by illuminating Batho at >520 nm for MB. This unique property of MB can be easily seen in Figure 2b, where the 9-*cis*-specific 1207 cm^−1^ bands were seen for MRh, MG, and MR, but not for MB.

### Y265 is Responsible for the Absence of the 9-cis Form in MB at 77 K

Animal rhodopsins generally exhibit photoequilibria of the 11-*cis*, all-*trans*, and 9-*cis* forms at 77 K.^3^ In contrast, the 9-*cis* form did not form in MB at 77 K, suggesting a unique chromophore binding pocket in MB. Figure 3 compares the 19 amino acids surrounding the retinal chromophore among MB, MG, MR, and MRh, based on the structure of bovine rhodopsin.^5^ It should be noted that W265 is highly conserved in animal rhodopsins including long (L group) and middle (M group) wavelength sensitive color visual pigments.^3,4^ In contrast, the corresponding residue of W265 is tyrosine in short wavelength sensitive color visual pigments (S group). Therefore, in this study, we mutated Y265 of MB into tryptophan, and W265 of MRh into tyrosine.

**Figure 3:**
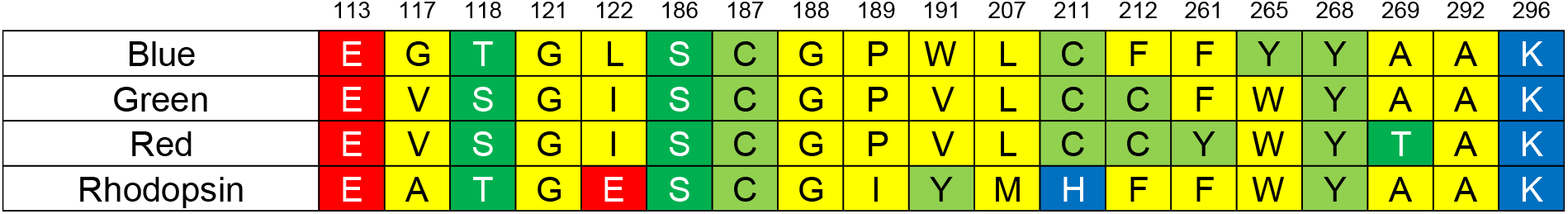
The amino acid residues surrounding the retinal chromophore in bovine rhodopsin (<4 A, PDB: 1U19),^5^ and the corresponding residues in MB, MG, MR and monkey rhodopsin.

Figure 4a clearly shows the dependence of illumination wavelength on Y265W MB, where dotted (400 nm and >520 nm lights) and solid (400 nm and >430 nm lights) lines differ significantly. The double difference spectrum in Figure 4b possesses a positive peak at 1201 cm^−1^, which has a different frequency from the 9-*cis*-specific band at 1207 cm^−1^ (Figure 3). Therefore, this is not a direct indication of the formation of the 9-*cis* state. Nevertheless, the present results clearly indicate a photoequilibrium of the three states, and it is reasonable to consider the 9-*cis* form as the third state in addition to the 11-*cis* (unphotolyzed) and all-*trans* (Batho) states. In this case, the peak frequency at 1201 cm^−1^ suggests that the 9-*cis* chromophore is not in a relaxed conformation. Another possibility is that the third state contains the 11-*cis* chromophore, whose absorption is blue-shifted from the fully relaxed 11-*cis* form. From the present observation, we conclude that the residue at position 265 contributes to the unique property of MB.

**Figure 4:**
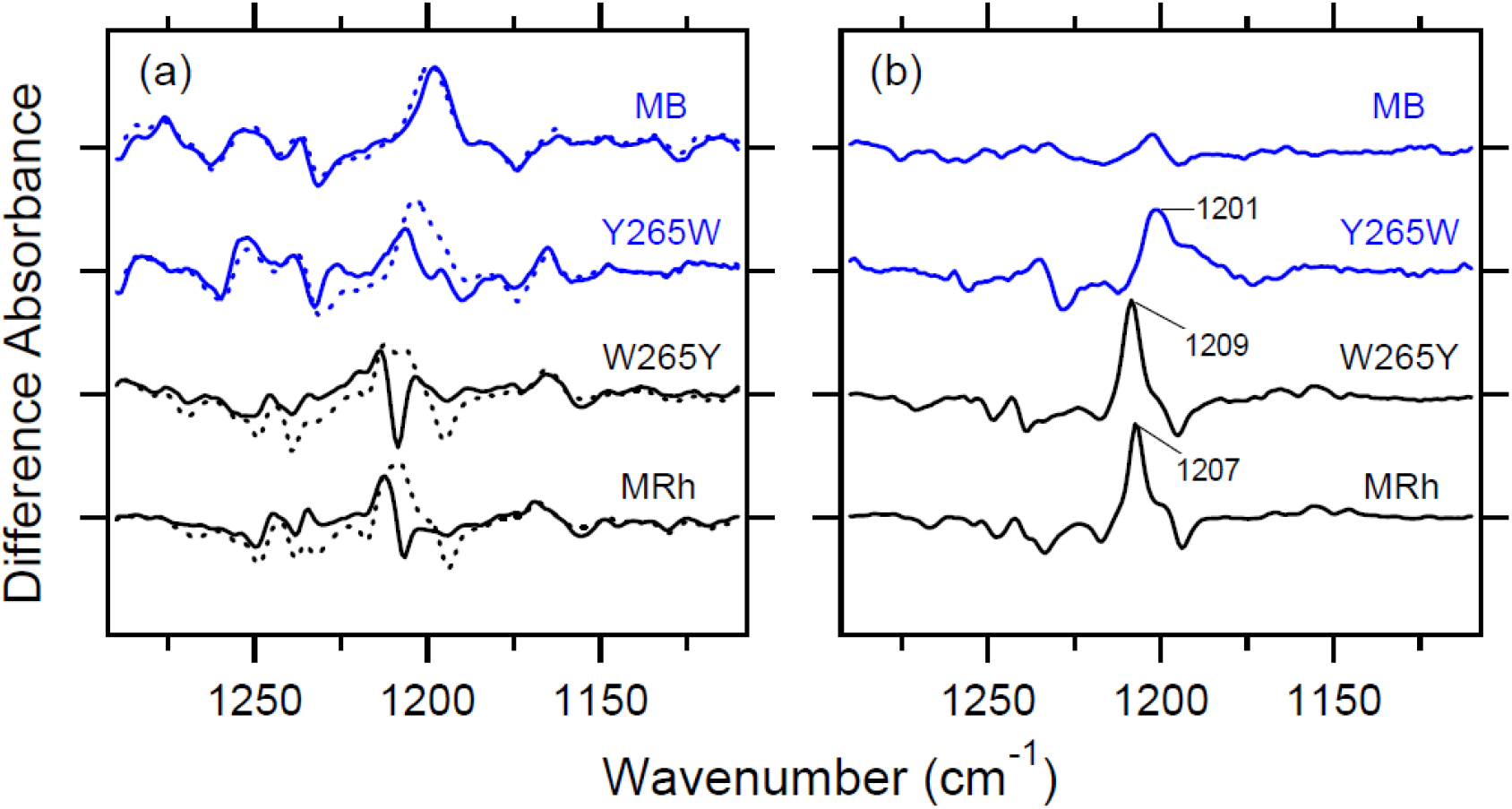
(a) Light-minus-dark difference FTIR spectra in the 1300-1100 cm^−1^ region for MB, Y265W MB, W265Y MRh, and MRh at 77 K. Repeated illumination conditions for dotted lines are 400 nm and >520 nm lights for MB and Y265W MB (blue line), and 501 nm and >610 nm lights for W265Y MRh and MRh (black line). Repeated illumination conditions for solid lines are 400 nm and >430 nm lights for MB and Y265W MB (blue line), and 501 nm and >540 nm lights for W265Y MRh and MRh (black line). (b) Double difference spectra in (a), where the solid line was subtracted from the dotted line.

In the case of MRh, the W265Y mutant exhibited similar spectral features to those of the wild-type protein (Figure 4a). In fact, a clear positive peak appeared at 1209 cm^−1^ in the double difference spectrum (Figure 4b). No conversion of MRh to the MB-specific property by this mutation suggests that the amino acid at position 265 is not the only determinant to produce the 9-*cis* form.

### No Pigment Formation from 9-cis Retinal and MB Opsin

The lack of formation of the 9-*cis* pigment at 77 K raised another issue, whether pigment could form from 9-*cis* retinal and MB opsin at room temperature. To clarify this, we divided HEK cells expressing MB into two, to which either 11-*cis* or 9-*cis* retinal was added. Then, the regenerated sample was solubilized by 1% DDM and illuminated in the presence of 10 mM hydroxylamine. Figure 5 shows the results of hydroxylamine bleach for the wild-type proteins of MB (a) and MRh (b), where dotted and solid lines represent the results for 11-*cis* and 9-*cis* retinal, respectively. When 11-*cis* retinal (dotted lines) was added, MB and MRh formed, and their absorption maxima are located at 430 and 499 nm, respectively. On the other hand, the 9-*cis* pigment did not form for MB (a), whereas 9-*cis* rhodopsin (isorhodopsin) formed for MRh, whose absorption maximum is located at 485 nm. The present results imply that the retinal binding pocket of MB does not accommodate 9-*cis* retinal, unlike other visual pigments.

**Figure 5:**
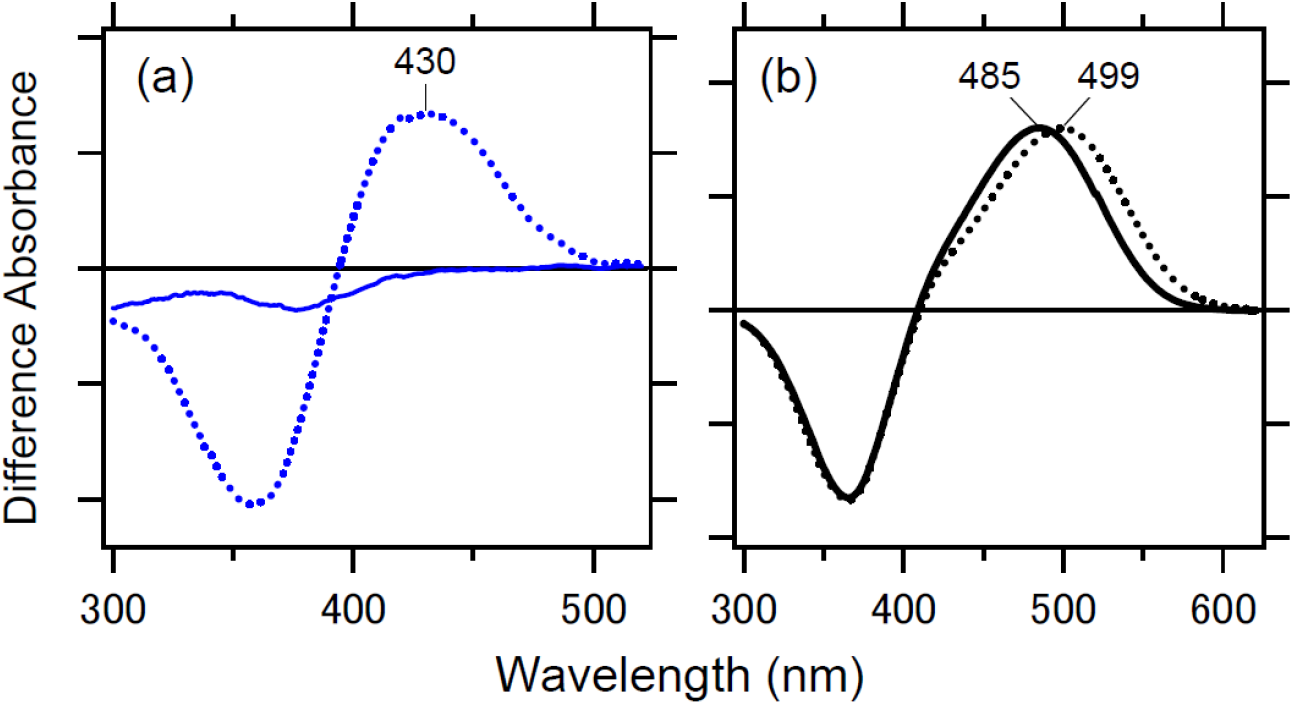
Difference UV-visible spectra of MB (blue line) and MRh (black line) by hydroxylamine bleach, where positive and negative absorption spectra originate from the unphotolyzed state and retinal oxime, respectively. Dotted and solid lines represent the results for 11-*cis* and 9-*cis* retinal, respectively. The DDM (final conc.: 1%)-solubilized proteins in the presence of 10 mM hydroxylamine were kept for 30 min in the dark at 4°C, and then illuminated with >400 nm (MB) or >480 nm (MRh) light for 1.5 min.

We also applied similar experiments to Y265W MB and W265Y MRh. These results, which are shown in Figure 6, were identical to the wild-type. Namely, retinal binding was observed for the 11-*cis* form of MB (dotted line in Figure 6a), and the 11-*cis* (dotted line in Figure 6b) and 9-*cis* (solid line in Figure 6b) forms of MRh, but not for 9-*cis* in MB (solid line in Figure 6a). Therefore, the Y265W mutation was not sufficient for pigment formation from 9-*cis* retinal and MB.

**Figure 6:**
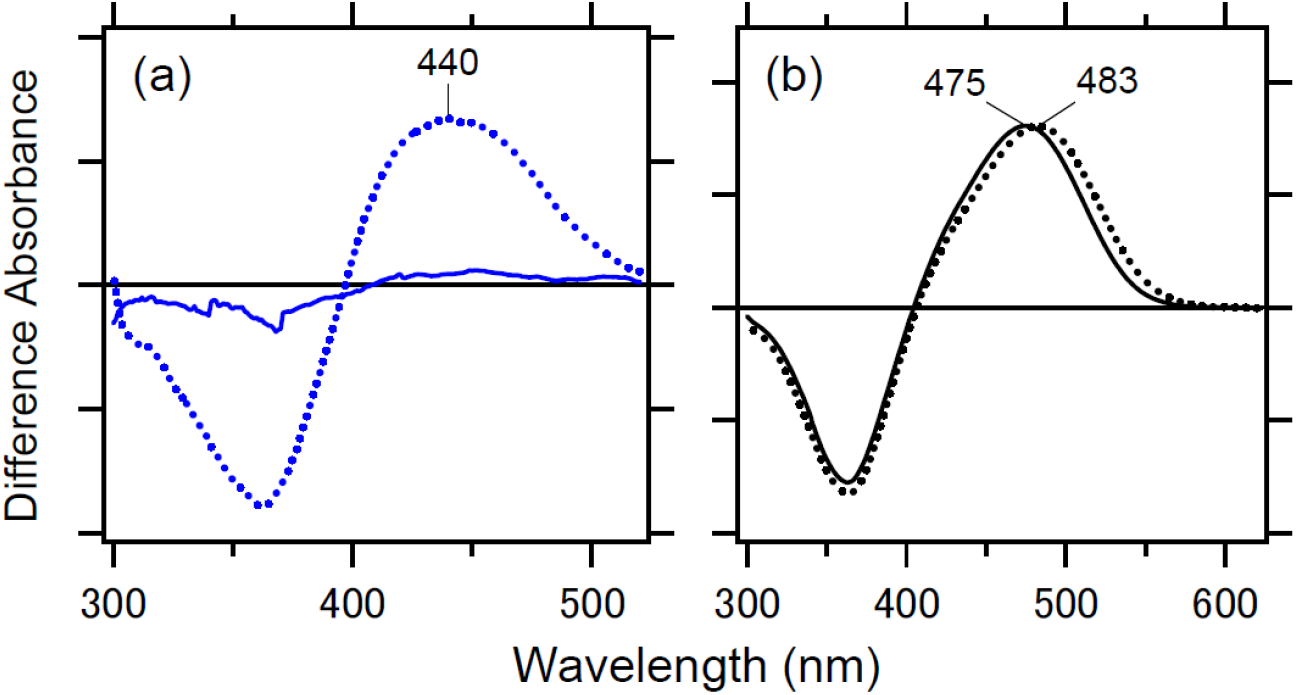
Difference UV-visible spectra of the mutant of position 265 for MB (Y265W MB; blue line) and MRh (W265Y MRh; black line) by hydroxylamine bleach, where positive and negative absorption spectra originate from the unphotolyzed state and retinal oxime, respectively. Dotted and solid lines represent the results for 11-*cis* and 9-*cis* retinal, respectively. The DDM (final conc.: 1%)-solubilized proteins in the presence of 10 mM hydroxylamine were kept for 30 min in the dark at 4°C, and then illuminated with >400 nm (Y265W MB) or >460 nm (W265Y MRh) light for 1.5 min.

## DISCUSSION

The retinal binding pocket is unique for visual pigments. In general, 11-*cis* retinal is not thermally stable and easily isomerizes into the all-*trans* form. On the other hand, 11-*cis* retinal is very stable in animal rhodopsins, indicating that the retinal binding pocket optimally accommodates the 11-*cis* form. Nevertheless, the 11-*cis* to all-*trans* photoisomerization takes places even at 77 K,^23,24^ where the protein environment is frozen. This is consistent with ultrafast retinal photoisomerization^36–40^ during which there is no time for the protein environment to change its structure. Note that the shapes of 11-*cis* and all-*trans* retinals differ, as shown in Figure 1. Therefore, the mechanism of retinal photoisomerization has been a long-standing issue in the study of vision.^3,41,42^

It is well known that the retinal binding pocket accommodates the 9-*cis* form in addition to the 11-*cis* and all-*trans* forms at 77 K. This is also the case for other visual pigments including MR and MG.^18^ However, the present study clearly demonstrates that the 9-*cis* form was not produced by illuminating 11-*cis* and all-*trans* forms in MB. It should be noted that the 9-*cis* form was produced for chicken blue-sensitive pigment at 77 K.^28^ Whereas chicken blue belongs to the M1-group, and chicken violet-sensitive pigment is in the S-group.^43^ A previous resonance Raman study attempted to answer the question “why are blue visual pigments blue?”, in which the 9-*cis* form was produced for the 440-nm absorbing pigment of the toad (*Bufo marinus*) at 77 K,^44^ although this pigment is not in the S-group. Thus we suggest that this property is unique to visual pigments in the S-group.

In the present study, we also showed that 9-*cis* retinal was not bound to MB opsin, which is also in striking contrast to other visual pigments. The lack of pigment formation from 9-*cis* retinal and human blue opsin was already reported in 1999, even though no data was shown.^45^ Therefore, it is likely that the unique retinal binding pocket is common for the S-group. Interestingly, it was revealed that 11-*cis*-locked 6-membered-ring retinal can be bound for human blue opsin, but neither for human green nor red opsin.^46^ These results also emphasize the uniqueness of the S-group.

These unique properties of the S-group originate from the amino acids that surround the retinal chromophore. The residues around the retinal chromophore based on the structure of bovine rhodopsin are shown in Figure 3, from which Y265 was found to play an important role in photoequilibrium at 77 K (Figure 4). A previous study with 11-*cis*-locked 6-membered-ring retinal also concluded that Y265 contributes significantly.^46^ The crystal structures of bovine rhodopsin^5^ and 9-*cis* rhodopsin^47^ in Figure 7 depict the unique position of W265, which is located at the bending region of the retinal chromophore. While the important role of position 265 is not in doubt, the present study also elucidated the contributions of positions other than position 265. Further experimental and theoretical studies are needed to clarify the unique chromophore-protein interaction in the S-group.

**Figure 7:**
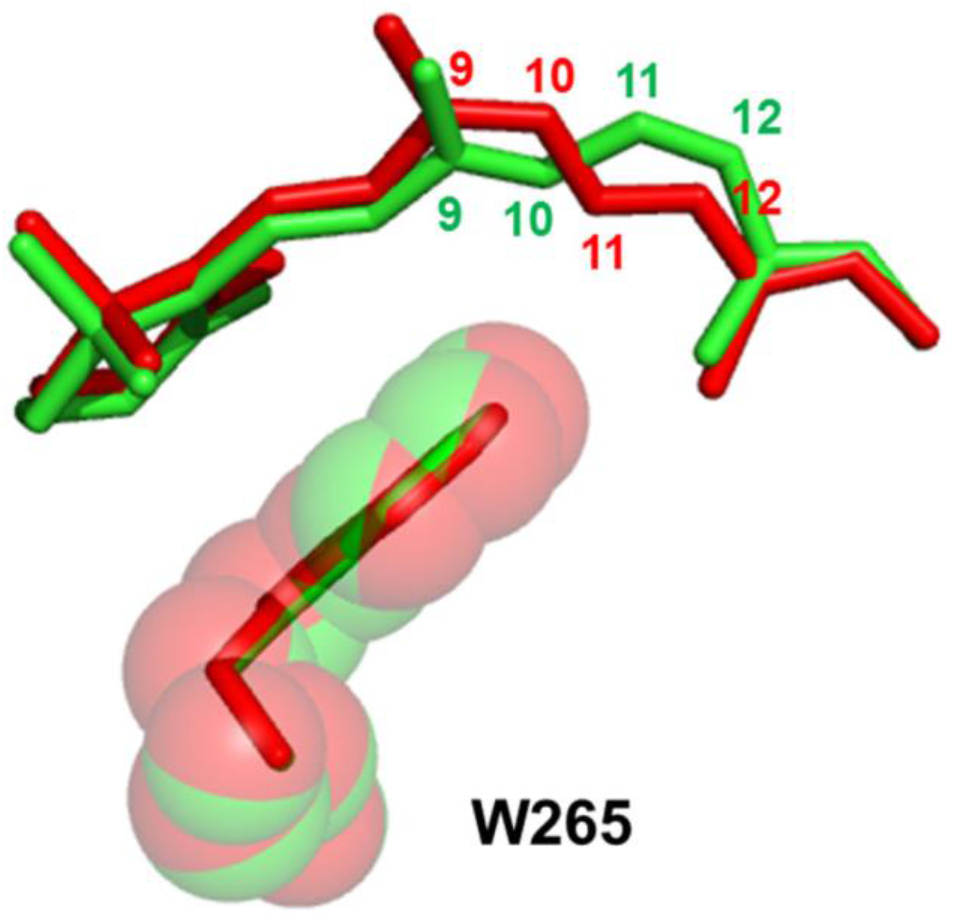
Structure of the chromophore and W265 in bovine rhodopsin containing 11-*cis* retinal (green; PDB: 1U19),^5^ and 9-*cis* retinal (red; PDB: 2PED).

## ACKNOWLEDGEMENTS

This work was financially supported by grants from the Japanese Ministry of Education, Culture, Sports, Science and Technology to K.K. (18K14662) and H.K. (18H03986, 19H04959), and from the Japan Science and Technology Agency (JST), PRESTO to K.K. (JPMJPR19G4). This work was partly supported by the JSPS Core-to-Core Program, A. Advanced Research Networks (Wildlife Research Center of Kyoto University).

## Conflicts of Interest

All authors declare no conflicts of interest.

## Notes

### Competing Interest Statement

The authors have declared no competing interest.

